# A palette of bright and photostable monomeric fluorescent proteins for bacterial time-lapse imaging

**DOI:** 10.1101/2024.03.28.587235

**Authors:** Nathan Fraikin, Agathe Couturier, Christian Lesterlin

## Abstract

Fluorescent proteins (FPs) are pivotal for examining protein production, localization, and dynamics in live bacterial cells. However, the use of FPs in time-lapse imaging is frequently constrained by issues such as oligomerization or limited photostability. Here, we report the engineering of four new fluorescent proteins: mChartreuse (green), mJuniper (cyan), mLemon (yellow), and mLychee (red), using site-directed mutagenesis. These fluorophores outperform current standards for photostability and aggregation properties while retaining high levels of brightness in living *E. coli* bacteria. Starting with superfolder GFP (sfGFP), we developed mChartreuse, which shows significantly enhanced brightness, stability, and monomericity compared to other GFP variants. From mChartreuse, we derived mJuniper, a cyan fluorescent protein with rapid maturation and high photostability, and mLemon, a yellow fluorescent protein with improved photostability and brightness. We also identified a mutation that eliminates residual oligomerization in red fluorescent proteins derived from *Discosoma* species, such as mCherry and mApple. Incorporating this mutation into mApple, along with other substitutions, resulted in mLychee, a bright and photostable monomeric red fluorescent protein. These novel fluorescent proteins advance fluorescence time-lapse analysis in bacteria and their spectral properties match current imaging standards, ensuring seamless integration into existing research workflows.

## Introduction

Fluorescent proteins (FPs) are essential tools in life science, enabling the tracking of protein localization and gene expression at the single-cell level (Chudakov et al., 2010). Green fluorescent protein (GFP) was initially cloned from the jellyfish *Aequorea victoria* and initiated a revolution in the study of protein localization (Chalfie et al., 1994). However, wild-type *A. victoria* GFP (avGFP) showed several shortcomings since it matured poorly at 37°C and required the use of phototoxic ultraviolet light (400 nm) to be excited (Ogawa et al., 1995). Mutagenesis of avGFP enabled the discovery of well-folded variants that were excited by blue light (488 nm), such as EGFP, GFPmut2, or superfolder GFP (sfGFP) (Cormack et al., 1996; Pédelacq et al., 2006; Zhang et al., 1996). GFP derivates also displayed a tendency to dimerize and cause mis-localization of fusion partners, which could be eliminated by disrupting a small hydrophobic dimerization interface with the A206K substitution (Zacharias et al., 2002). Substitution of residues critical for fluorescence also enabled the engineering of blue, cyan, and yellow variants of GFP, expanding the palette of usable FPs (Heim et al., 1994; Ormö et al., 1996). The discovery of a red fluorescent protein (RFP) in *Discosoma sp*. paved the way for multicolor analysis. Indeed, this *Discosoma sp*. red fluorescent protein (DsRed) was homologous to avGFP but excited by green light (558 nm), which therefore enabled simultaneous imaging of GFP and DsRed with minimal spectral overlap (Matz et al., 1999). However, engineering usable derivatives of DsRed was extremely challenging since this red FP matures slowly, oligomerizes as a tight tetramer, and can stochastically mature as a green side-product that overlaps with the spectral properties of GFP (Bevis and Glick, 2002). Bevis & Glick overcame the slow maturation of DsRed by random mutagenesis, yielding DsRed-Express, which matured 15 times faster than DsRed (Bevis and Glick, 2002). A *tour-de-force* study by the Tsien lab reported the monomerization of DsRed-Express into a fast-maturing monomeric RFP (mRFP1) through structure-guided disruption of its AB and AC tetramerization interfaces (Campbell et al., 2002). While mRFP1 was dim and bleached quickly when excited, subsequent studies evolved a palette of DsRed derivatives with a wide range of properties, which were called mFruits (Shaner et al., 2008, 2004). Of all mFruits, mCherry became the standard RFP due to its fast maturation, high photostability, and lack of GFP-like side product (Shaner et al., 2004).

Over the years, potentially useful fluorescent proteins were identified in other organisms. For example, an extremely bright yellow FP was identified in the cephalochordate *Branchiostoma lanceolatum* and engineered in a monomeric green FP called mNeonGreen (mNG) (Shaner et al., 2013). mNeonGreen was twice as bright as EGFP, with similar photostability and maturation kinetics as the latter (Shaner et al., 2013), thereby highlighting the potential for bright and exploitable FPs to be discovered in organisms beyond *Discosoma sp*. or *A. victoria*. Other recently released FPs such as AausFP1 from *Aequorea australis*, mStayGold from *Cyteais uchidae* or AzaleaB5 from *Montipora monasteriata* show great promise as bright fluorescent probes (Ando et al., 2023b, 2023a; Lambert et al., 2020). Novel FPs could also be engineered *de novo* from a synthetic template as was the case for the mScarlet family (Bindels et al., 2016). Further evolution of mScarlet yielded mScarlet3 and mScarlet-I3, the brightest red FPs to date (Gadella et al., 2023).

All FPs have different sets of parameters that dictate setups for which they should be used. These include fluorescence spectrum, brightness, maturation rate, photostability and dispersibility (Cranfill et al., 2016). The brightness of FPs is measured as the product of product of its extinction coefficient and quantum yield and represents the amount of signal that can be obtained from a given FP. However, another critical parameter that influences *in vivo* brightness is maturation rate, which defines the kinetics by which a folded dark FP is converted into a fluorescent species by cyclization of three amino acids that become its chromophore. Because bacteria tend to be fast-growing organisms, the use of fast-maturing FPs is of utmost importance in these organisms to achieve high brightness, since dark species of slow-maturing FPs would quickly get diluted by fast doublings (Balleza et al., 2018).

Here, we used site-directed mutagenesis to engineer high-performance fluorescent proteins. We report the development of four new FPs that offer superior brightness and photostability while being truly monomeric. We developed mChartreuse from sfGFP by introducing 6 mutations (N39I, I128S, D129G, F145Y, N149K & V206K) that enhanced brightness, photostability and monomericity (**Figure 1**). We further developed mChartreuse into mJuniper, a cyan variant, with the addition of four mutations (Y66W, S72A, N146F & H148D), and into mLemon, a yellow variant, with the addition of five mutations (T63S, T65G, S72A, T203Y & V224L) (**Figure 1**). Compared to state-of-the-art cyan and yellow FPs, these two FPs offer superior dispersibility and photostability with similar brightness (**Table 1**). While engineering red FPs, we identified S131P, a substitution that eliminates oligomerization of supposedly monomeric DsRed derivatives in *E. coli*. By combining S131P with brightness-enhancing mutations (V71A, L85Q, K139R, A145P & I210V), we evolved mApple into mLychee, a bright, monomeric and photostable red FP (**Figure 1, Table 1**). We therefore offer a palette of novel high-performance fluorescent proteins tested in *E. coli*, developed by microbiologists for microbiologists.

**Table 1:** Properties of purified fluorescent proteins. Columns show means (± standard deviation when applicable) with units in parenthesis: Fluorescence protein (FP), Absorbance (Abs) peak, Emission (Em) peak, Extinction coefficient (EC), Quantum yield (QY), Brightness (product of EC & QY), pKa & maturation half-time.

| FP | Abs peak (nm) | Em peak (nm) | EC (mM<sup>-1</sup> cm<sup>-1</sup>) | QY | Brightness | pKa | Maturation (min) |
| --- | --- | --- | --- | --- | --- | --- | --- |
| mChartreuse | 487 | 510 | 71±3 | 0.75±0.02 | 53±3 | 4.9±0.3 | 4.9±0.6 |
| mJuniper | 434 | 475 | 32±1 | 0.43±0.04 | 14±2 | 4.7±0.1 | 1.7±1.1 |
| mLemon | 514 | 526 | 125±5 | 0.74±0.03 | 93±6 | 4.8±0.1 | 12.0±1.1 |
| mLychee | 568 | 596 | 80±4 | 0.50±0.02 | 40±3 | 6.3±0.1 | 36.4±2.2 |

**Figure 1:**
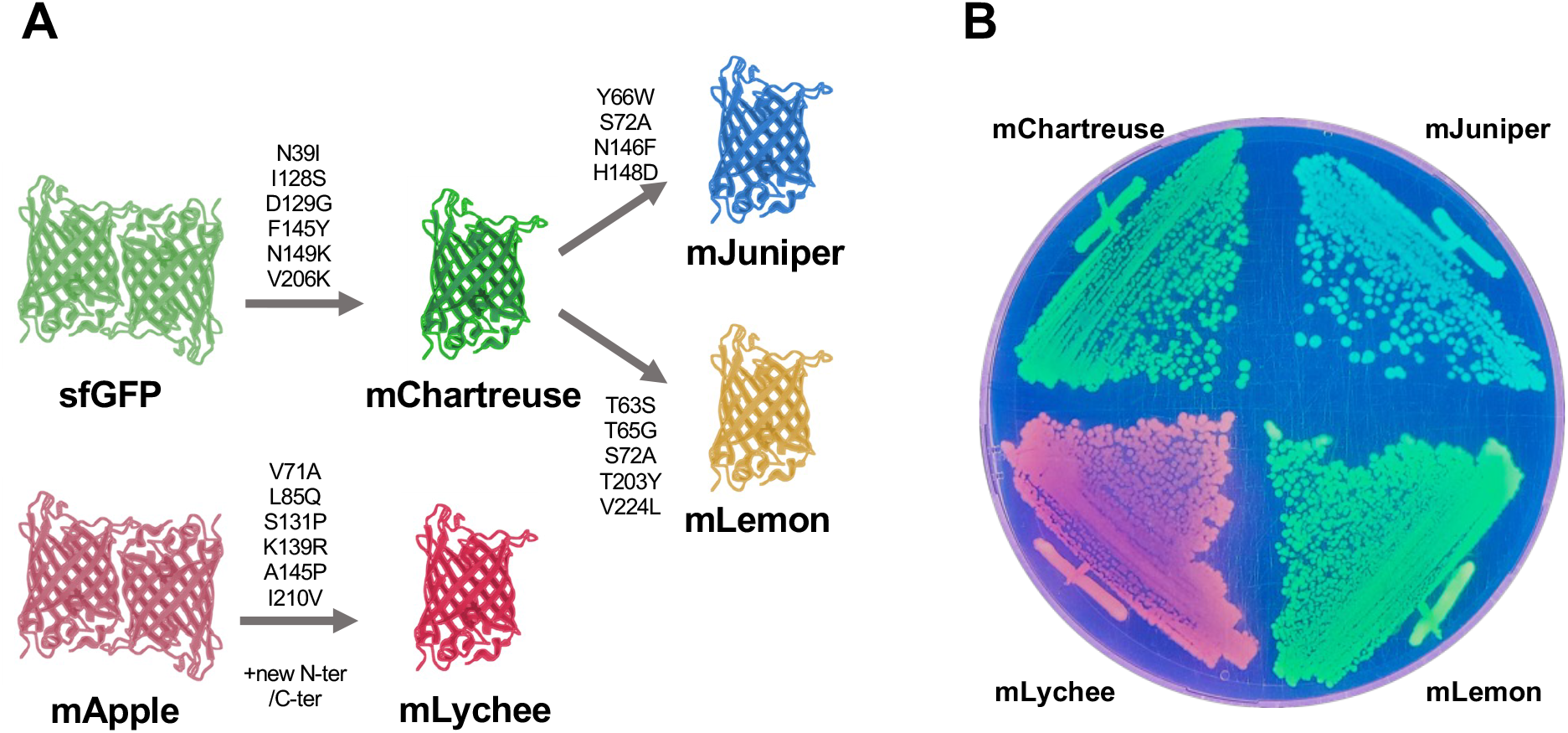
Novel fluorescent proteins detailed in this study. **A:** Flow chart of fluorescent proteins developed in this study. Each arrow shows a development step with its associated substitutions. New N-ter/C-ter refers to the replacement of mApple N and C-termini with those of mCherry2C (Valbuena et al., 2020). **B:** Images of bacterial colonies carrying pNF02 vectors encoding each indicated fluorescent proteins, trans-illuminated with UV light.

## Results

### Site-directed mutagenesis improves all properties of sfGFP

We first aimed to obtain an improved GFP derived from the well-characterized *A. victoria* GFP (avGFP) scaffold. We chose sfGFP as a starting template due to its improved folding, fast maturation, high brightness and broad use in bacterial models (Balleza et al., 2018; Pédelacq et al., 2006). We first introduced the V206K monomerizing mutation and the F145Y photostability-improving mutation as previously described for the construction of msGFP2 (Valbuena et al., 2020). To further improve the brightness of this starting template, we used high-throughput flow cytometry to screen several mutations found in bright avGFP derivatives. After four rounds of site-directed mutagenesis, we identified our brightest variant containing mutations N39I, I128S, D129G from the Achilles YFP (Yoshioka-Kobayashi et al., 2020) and N149K from the Emerald GFP (Teerawanichpan et al., 2007) (**Figure 2A, Supplementary Figure 1**). We called this variant mChartreuse based on the similarly colored liquor from southeast France.

**Figure 2:**
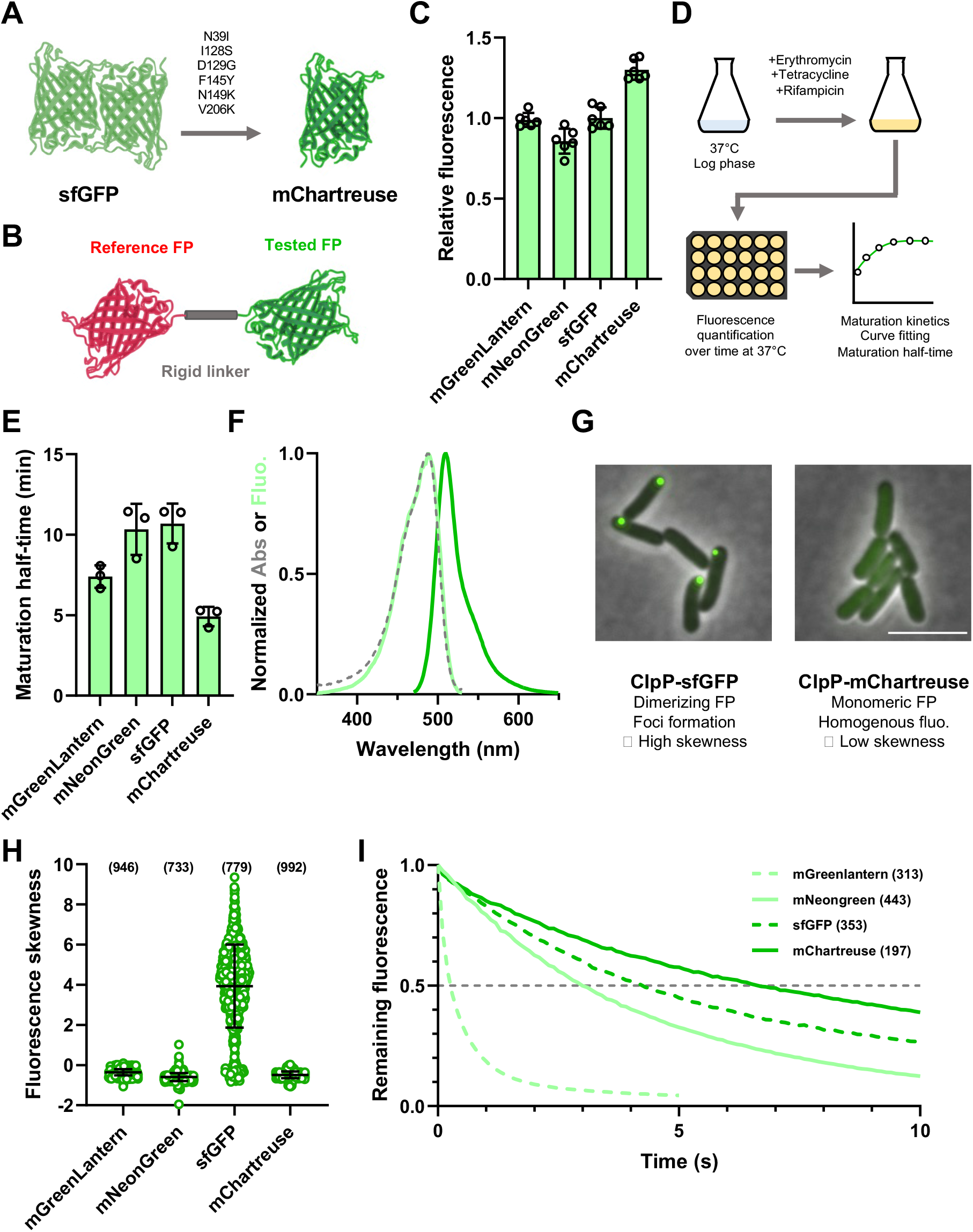
Development and characterization of the mChartreuse green fluorescent protein. **A:** All six mutations introduced in sfGFP to obtain mChartreuse. **B:** Representation of the brightness quantification system developed by (Bindels et al., 2016), where the tested FP is fused to a spectrally distinct FP. The ratiometric measurement of the tested FP normalized to that of the reference FP enables to offset any variation in FP signal due to differences in expression levels. The rigid spatial linker that separates both FPs prevents fluorescence resonance energy transfer (FRET). **C:** *In vivo* brightness quantification of green FPs.

Green FP fluorescence was measured in exponentially growing cultures and normalized to that of mCherry. Data shows mean and standard deviation of six independent replicates. **D:** Description of the maturation assay. Exponentially growing cells were treated with 100 µg/mL erythromycin, 10 µg/mL tetracycline hydrochloride & 10 µg/mL rifampicin to inhibit protein synthesis and enable full maturation of dark u nmatured fluorophores. Maturation kinetics were measured from this point and fitted to a pseudo-first order kinetics, from which maturation half-times were computed. **E:** Maturation half-times of green FPs. Data shows mean and standard deviation of three independent replicates. **F:** Spectral properties of mChartreuse. The dashed line shows the absorbance spectrum, the light-colored line shows the excitation spectrum, and the dark-colored line shows the emission spectrum. Data is shown as the mean of three independent spectrum acquisitions. **G:** Representative images of ClpP fusions. Fusion of ClpP with a dimeric FP (*i*.*e*. sfGFP) causes substantial aggregation, which leads to the formation of fluorescence foci and high fluorescence skewness in single cells. Fusion with a monomeric FP (*i*.*e*. mChartreuse) does not cause aggregation, leading to homogenous FP distribution in the cytosol and low fluorescence skewness. **H:** Fluorescence distribution skewness of single cells expressing ClpP-FP fusions. Bars show the mean and standard deviation. Data were acquired from three independent experiments, with the total numbers of analyzed bacteria shown in brackets. **I:** Photostability of FP signals in live bacteria illuminated with 100 ms steps and normalized to intensity at time 0. The dashed grey line shows the cutoff at which t_1/2_ were calculated. Data were acquired from three independent experiments, with the total numbers of analyzed bacteria shown in brackets.

Purified mChartreuse displayed an extinction coefficient of 71 mM^-1^cm^-1^. and a quantum yield of 0.75 (**Table 1**). However, evaluating *in vivo* brightness is a better indicator of FP performance in living systems. By fusing mChartreuse and other GFPs with mCherry, we can evaluate the *in vivo* brightness of these FPs in growing cells by normalizing the green fluorescence signal to that of mCherry, therefore allowing us to offset any variation in the expression level of these constructs (Bindels et al., 2016) (**Figure 2B**). Our results show that mChartreuse is 30% brighter than sfGFP in *E. coli* (**Figure 2C**). mChartreuse is also brighter than mNeonGreen and mGreenLantern, two bright and top-performing GFPs that will be used hereafter as comparison points to sfGFP and mChartreuse (Campbell et al., 2020; Shaner et al., 2013) (**Figure 2C**).

To assess how well mChartreuse matured in exponentially growing *E. coli*, we quantified the maturation half-time of mChartreuse and other green FPs by shutting down protein synthesis using a cocktail of erythromycin, tetracycline and rifampicin, followed by measuring fluorescence maturation kinetics after protein synthesis shutoff (**Figure 2D**). In this setup, protein synthesis inhibition arrests fluorophore biosynthesis, allowing remaining dark unmatured fluorophores to fully mature (**Supplementary Figure 2A**). By fitting a pseudo-first order kinetics curve to the maturation data, we determined the maturation half-time of mChartreuse to be 4.9 min, faster than its sfGFP parent (10.7 min), or than mNeonGreen (10.3 min) and mGreenLantern (7.4 min) (**Table 1, Figure 2E)**. The excitation and emission spectra of untagged mChartreuse were determined after three phase partition purification (Jain et al., 2004). mChartreuse displayed a broad absorbance peak with a maximum at 487 nm, while its emission peaked at 510 nm, similar to that of superfolder GFP or EGFP (Pédelacq et al., 2006; Zhang et al., 1996) (**Table 1, Figure 2F**). The fluorescence of mChartreuse displays a pKa of 4.9, showing that it is resistant to fluctuations in cytosolic pH (**Table 1, Supplementary Figure 3**)

We evaluated the dispersibility and oligomerization of mChartreuse and other green FPs by fusing them to the ClpP protease as previously described (Landgraf et al., 2012). In this system, oligomeric FPs cause cooperative clustering of ClpP homo-tetradecamers into fluorescent foci, while monomeric FPs do not perturbate the homogenous cytosolic localization of ClpP (Landgraf et al., 2012). To offer an unbiased quantification of foci formation by these ClpP fusions, we measured the skewness of pixel intensity distributions in single bacteria as a proxy for fluorescence inhomogeneity (Ducret et al., 2016). As previously described, fusion of ClpP with sfGFP (a dimerizing FP) induced foci formation, exemplified by the high fluorescence skewness observed in such cells (**Figure 2G-H**). On the other hand, ClpP fusions with mChartreuse, mNeonGreen, and mGreenLantern all displayed homogeneous fluorescence as shown by the low fluorescence skewness values of these fusions (**Figure 2G-H**). Our results therefore show that mChartreuse is monomeric in *E. coli*.

Because photostability is a critical parameter for time-lapse imaging, we evaluated the time required to bleach mChartreuse to half its original fluorescence intensity (t_1/2_) by time-lapse fluorescence microscopy. Of tested GFPs, mChartreuse displayed the highest photostability (t_1/2_ = 6.7 s), followed by sfGFP (t_1/2_ = 4.3 s), mNeonGreen (t_1/2_ = 3.0 s), and finally mGreenLantern (t_1/2_ = 0.3 s) (**Figure 2I**). Conclusively, the evolution of mChartreuse improved all aspects of sfGFP (*i*.*e* brightness, maturation, dispersibility and photostability), making it a superior choice for all applications.

### Site-directed mutagenesis of mChartreuse yields highly photostable monomeric cyan and yellow FPs

Since mChartreuse is a bright and photostable variant of sfGFP, we assessed whether this FP could be engineered into cyan and yellow variants with superior properties. We developed a cyan variant of mChartreuse by introducing the Y66W cyan-determining mutation (Heim et al., 1994), followed by the S72A, N146F and H148D mutations found in mTurquoise2 (Goedhart et al., 2012) (**Figure 3A, Supplementary Figure 1**). This variant, named “mJuniper”, was compared to SCFP3A and mTurquoise2, two top-performing CFPs (Goedhart et al., 2012; Kremers et al., 2006). The brightness of mCherry-fused mJuniper was equivalent to that of SCFP3A, with both these FPs being roughly 50% brighter mTurquoise2 (**Figure 3B**). mJuniper displayed unusually fast maturation kinetics, with a maturation half-time of 1.7 min, faster than SCFP3A (6.9 min) or mTurquoise2 (36.5 min) (**Table 1, Figure 3C, Supplementary Figure 2B**). mJuniper showed an absorbance peak at 434 nm, and an emission peak at 475 nm as described for SCFP3A and mTurquoise2 (Goedhart et al., 2012; Kremers et al., 2006) (**Table 1, Figure 3D**). The pKa of mJuniper is 4.7, showing that it can withstands changes in cytosolic pH (**Table 1, Supplementary Figure 3**). Fusions of ClpP with mTurquoise2 and SCFP3A both lead to foci formation in a significant number of cells while fusion of ClpP with mJuniper yielded homogenous fluorescence, showing that mJuniper is a true monomer (**Figure 3E**). Moreover, mJuniper displayed higher photostability (t_1/2_ = 2.2 s) compared to mTurquoise2 (t_1/2_ = 1.1 s) and SCFP3A (t_1/2_ = 1.1 s) (**Figure 3F**). Therefore, mJuniper is a bright and truly monomeric cyan FP with superior maturation and photostability in *E. coli*.

**Figure 3:**
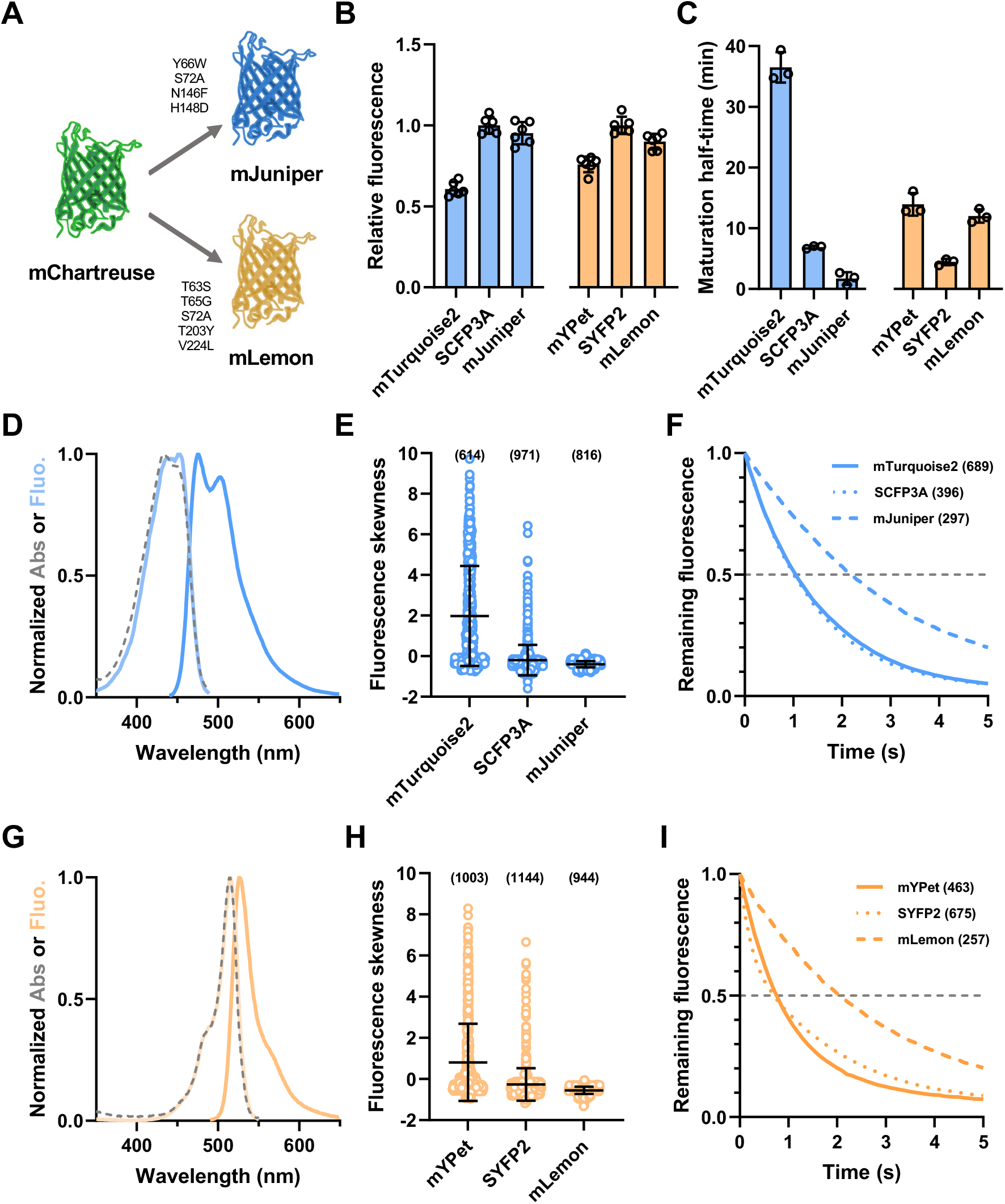
Development and characterization of cyan and yellow variants of mChartreuse. **A:** Mutations introduced in mChartreuse to obtain the mJuniper cyan variant and the mLemon yellow variant. **B:** *In vivo* brightness quantification of cyan and yellow FPs. Fluorescence was measured in exponentially growing cultures and normalized to that of mCherry (for cyan FPs) or mTurquoise2 (for yellow FPs). Data shows mean and standard deviation of six independent replicates. **C:** Maturation half-times of cyan and yellow FPs. Data shows mean and standard deviation of three independent replicates. **D & G:** Spectral properties of mJuniper (**D**) and mLemon (**G**). The dashed line shows the absorbance spectrum, the light-colored line shows the excitation spectrum and the dark-colored line shows the emission spectrum. Data is shown as the mean of three independent spectrum acquisitions. **E & H:** Fluorescence distribution skewness of single cells expressing ClpP-CFP (**E**) or ClpP-YFP (**H**) fusions. Bars show the mean and standard deviation. Data were acquired from three independent experiments, with the total numbers of analyzed bacteria shown in brackets. **F & I:** Photostability of CFP signals (**F**) and YFP signals (**I**) in live bacteria illuminated with 100 ms steps and normalized to intensity at time 0. The dashed grey line shows the cutoff at which t_1/2_ were calculated. Data were acquired from three independent experiments, with the total numbers of analyzed bacteria shown in brackets.

A yellow variant of mChartreuse was engineered by first introducing the T203Y yellow-determining mutation, followed by the T65G and S72A found in most yellow derivatives of avGFP (Ormö et al., 1996) (**Figure 3A, Supplementary Figure 1**). We also added the T63S mutation from mGold and the V224L from YPet (Lee et al., 2020; Nguyen and Daugherty, 2005) (**Figure 3A**). We called this variant “mLemon” and compared it to mYPet (YPet A206K) and SYFP2 (mVenus L68V), two top-performing YFPs (Kremers et al., 2006; Nguyen and Daugherty, 2005). SYFP2 and mLemon were both brighter than mYPet, with mLemon showing 90% of the brightness of SYFP2 (**Figure 3B**). mLemon matured at a similar rate as mYPet (12.0 min & 13.9 min, respectively), both maturing slower than SYFP2 (4.4 min) (**Table 1, Figure 3C, Supplementary Figure 2C**). The spectrum of mLemon was similar to that of other yellow FPs (Kremers et al., 2006; Nguyen and Daugherty, 2005), with narrow absorbance and emission peaks at 514 nm and 526 nm, respectively (**Table 1, Figure 3G**). mLemon showed high resistance to acid with a pKa of 4.7 (**Table 1, Supplementary Figure 3**). While some yellow FPs can be quenched by chloride ions (Wachter and James Remington, 1999), we show mLemon fluorescence to be insensitive to increases in sodium chloride concentration (**Supplementary Figure 4**). ClpP-mLemon displayed homogenous fluorescence while ClpP-mYPet and ClpP-SYFP2 both showed aggregation, showing that mLemon is monomeric (**Figure 3H**). mLemon displayed more than twice the photostability (t_1/2_ = 2.1 s) of mYPet (t_1/2_ = 0.8 s) and SYFP2 (t_1/2_ = 0.8 s) (**Figure 3I**). Therefore, mLemon offers higher photostability and lower aggregation compared to other yellow FPs while still retaining high brightness.

### Monomerization and enhancement of the mApple red FP

We also sought to engineer a bright and photostable monomeric red fluorescent protein. While mScarlet-I3 was monomeric and offered unrivaled brightness, it showed lower photostability compared to other red FPs (Gadella et al., 2023). On the other hand, mFruits like mCherry and mApple offered higher photostability (Shaner et al., 2008, 2004). However, previous data shows that mFruits aggregate when fused to ClpP in *E. coli* (Landgraf et al., 2012). Indeed, our data shows that fusions of mCherry and mApple with ClpP had a high propensity to form foci, confirming that these FPs are not monomeric (**Figure 4A**). On the other hand, ClpP-mScarlet-I3 did not form foci, indicating that this FP is truly monomeric (**Figure 4A**).Since mScarlet-I3 was derived from a mFruit-based synthetic template, we introduced mutations specific to mScarlet-I3 into mApple, then screened for mutations conferring a non-aggregative behavior in ClpP fusions. The S131P mutation (numbering relative to DsRed) abolished foci formation in ClpP fusions with mApple or mCherry (**Figure 4A**). In DsRed tetramers,S131 is part of the AB tetramer interface between two DsRed protomers, with the hydroxyl sidechain and the amide nitrogen of S131 each implicated in a hydrogen bond with the carboxyl sidechain of D154 located *in trans* (**Figure 4B**). It is therefore likely that the substitution of S131 with a proline prevents H-bond formation with D154 and eliminates residual dimerization of mFruits in *E. coli*. we used mApple-S131P as a starting template to engineer a brighter RFP. During the previous engineering of mFruits, the original N and C-termini of DsRed were replaced with those of avGFP (Shaner et al., 2004). While this allowed proper localization of some fusions, restoration of DsRed-like termini was shown to improve brightness and relieve RFP-induced cytotoxicity (Gadella et al., 2023; Valbuena et al., 2020). We therefore synthetized a codon-optimized mApple template, replaced the MVSKGEEN NM N-terminus and TGGMDE C-terminus of mApple with the MDSTE N-terminus and GSQGGSGGS C-terminus of mCherry2C (**Supplementary Figure 5**) (Valbuena et al., 2020). We subsequently performed rounds of site-directed mutagenesis to identify mutations conferring increased *in vivo* brightness. Mutations V71A, L85Q, K139R, A145P and I210V were incorporated in our final variant, baptized mLychee (**Figure 4C,Supplementary Figure 5**).

The mLychee red FP is 28% brighter than mApple, and six times brighter than mCherry under excitation at 565 nm, while mScarlet-I3 outperforms other FPs, being more than thrice as bright as mApple (**Figure 4D**). mLychee showed a maturation half-time of 36.4 min, faster than its mApple parent (55.0 min) or than mCherry (44.4 min), but slower than mScarlet-I3 (21.5 min) (**Table 1, Figure 4E,Supplementary Figure 2D**).Similarly to what was published for mScarlets and mApple (Bindels et al., 2016; Gadella et al., 2023;Shaner et al., 2008), mLychee displays an absorbance peak at 568 nm, an emission peak at 596 nm and a pKa of 6.3 (**Table 1, Figure 4F,Supplementary Figure 3**). mCherry displayed the highest photostability (t_1/2_ = 3.6 s) while mScarlet-I3 bleached the quickest (t_1/2_ = 0.4 s) (**Figure 4G**). mApple and mLychee displayed complex bleaching behaviors, with an abrupt loss of 15% of their signal after the first 100 ms exposure, followed by slow bleaching kinetics similar to that of mCherry (t_1/2_ = 2.7 s & 2.8 s, respectively) (**Figure 4G**). Previous results showed that this fast initial bleaching of mApple was a photochromic phenomenon reversible either by time or by illuminating this fluorophore with blue light (Shaner et al., 2008). We therefore quantified the bleaching of mLychee by alternating illuminations between 555 nm and 474 nm (**Figure 4H**). In these conditions, the fast initial bleaching of mLychee was practically eliminated, yielding initial bleaching kinetics similar to that of mCherry (**Figure 4H**). Therefore, mLychee is a truly monomeric red FP that offers an excellent balance between brightness and photostability.

**Figure 4:**
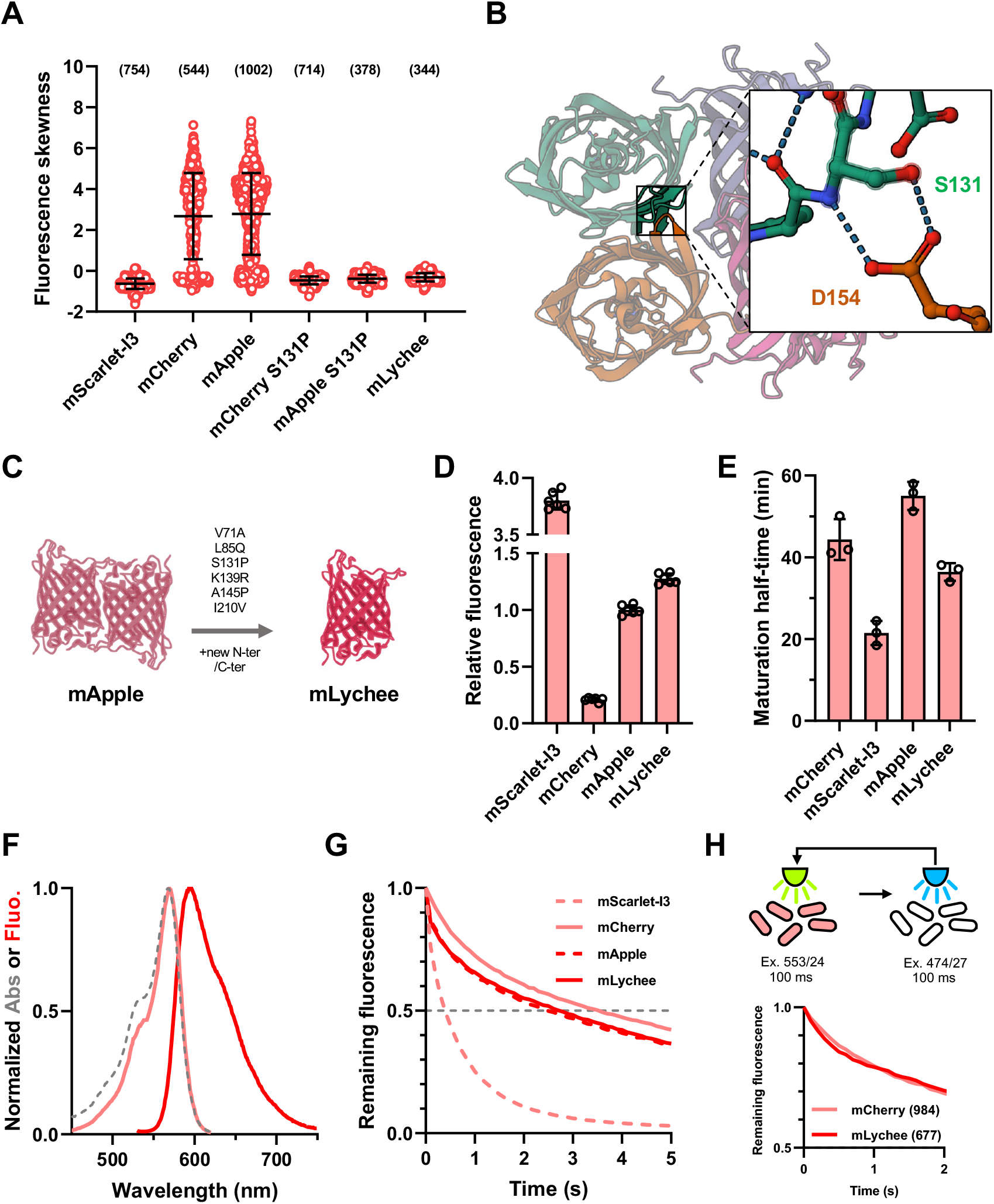
Development and characterization of the mLychee red fluorescent protein. **A:** Fluorescence distribution skewness of single cells expressing ClpP-FP fusions. Bars show the mean and standard deviation. Data were acquired from three independent experiments, with the total numbers of analyzed bacteria shown in brackets. **B:** Location of serine 131 (S131) in the AB dimerization interface of a DsRed tetramer (PDB 1ZGO)shown in ribbon representation. The inlet shows the interaction betweenS131 and aspartate 154 (D154) located *in trans* at the atomic level, with putative hydrogen bonds shown as dashed lines. **C:** All six mutations introduced in mApple to obtain mLychee. **D:** *In vivo* brightness quantification of red FPs. Fluorescence was measured in exponentially growing cultures and normalized to that mTurquoise2. Data shows mean and standard deviation of six independent replicates. **E:** Maturation half-times of red FPs. Data shows mean and standard deviation of three independent replicates. **F:**Spectral properties mLychee. The dashed line shows the absorbance spectrum, the light-colored line shows the excitation spectrum and the dark-colored line shows the emission spectrum. Data is shown as the mean of three independent spectrum acquisitions. **G:** Photostability of FP signals in live bacteria illuminated with 100 ms steps and normalized to intensity at time 0. The dashed grey line shows the cutoff at which t_1/2_ were calculated. Data were acquired from three independent experiments, with the total numbers of analyzed bacteria shown in brackets. **H:**Short-term photostability of mCherry and mLychee signals under alternating green (555 nm) and blue (474 nm) illumination. Data were acquired from three independent experiments, with the total numbers of analyzed bacteria shown in brackets.

## Discussion

In this manuscript, we introduce a novel palette of fluorescent proteins tailored for bacterial applications. This toolkit that includes mChartreuse, mJuniper, mLemon, and mLychee, represents a significant advancement in time-lapse imaging with its improved properties compared to traditional and commonly used fluorescent proteins. The enhanced photostability and minimal aggregation of these novel proteins make them ideal fusion tags, and their spectral characteristics enable straight forward integration into existing research workflows without requiring specialized equipment. Contrasting with other modern bright FPs like mNeonGreen or StayGold, mChartreuse and its derivatives were engineered from sfGFP, a well-folded green FP that is broadly described and used in bacterial imaging studies. Applications that currently use sfGFP can therefore be quickly adapted to use mChartreuse, mJuniper and mLemon. Further testing of these novel FPs as fusion tags in *E. coli* and other bacterial species will enable us to assess how well such fusions preserve native localization and activities of essential prokaryotic proteins.

Our findings confirm that the F145Y mutation significantly improves the photostability of sfGFP derivatives, consistent with previous research (Valbuena et al., 2020). However, the relationship between oligomerization and residue substitution appears more intricate. To examine FP oligomerization in *E. coli*, we developed a quantitative method based on the skewness in fluorescence distribution of ClpP fusions within single bacteria, enabling unbiased and robust quantification of FP aggregation and foci formation. Our results confirm that the V206K mutation abolishes oligomerization in sfGFP derivatives (Valbuena et al., 2020). However,since the A206K mutation is also present in mVenus and mYpet, which tend to aggregate in the ClpP assay (Landgraf et al., 2012), this suggests that aggregation behavior is influenced by factors beyond this mutation,such as superfolder mutations present in our initial sfGFP template (S30R, F99S, N105T, I171V) or mutations that were introduced during the development of mChartreuse (N39I, I128S, D129G, F145Y, N149K).

Previous research showed that mCherry and mApple are monomers in mammalian cells when assayed using the Organized Smooth Endoplasmic Reticulum (OSER) approach (Bindels et al., 2016; Cranfill et al., 2016). However, fusions of mCherry with ClpP, IbpA or RpoS in *E. coli* all caused substantial aggregation (Goormaghtigh et al., 2018; Govers et al., 2018; Landgraf et al., 2012),suggesting that red fluorescent proteins derived from *Discosoma sp*, are not monomeric in bacterial cells. The reasons for these discrepancies are not known but might stem from differences in cytosolic composition and crowding between mammalian and bacterial cells, or in differences of sensitivity between the OSER and ClpP assays. Nevertheless, the substitution of the serine 131 of mApple by a proline abolishes the aggregating behavior of the mLychee variant developed from mApple. While mLychee is dimmer than mScarlet-I3, it compensates for this lack of brightness with robust photostability. mLychee would therefore be more appropriate for time-lapse applications while mScarlet-I3 would remain preferable for endpoint methods (*e*.*g*., epifluorescence microscopy snapshots, flow cytometry) where photobleaching due to repeated excitation is not an issue.

Although these novel fluorescent proteins were initially developed for bacterial applications, they have the potential to significantly enhance imaging techniques in eukaryotic cells as well. One major consideration in transferring these FPs to eukaryotic systems is their compatibility with the cellular environment, including cytosolic composition, protein expression, and folding dynamics, which differ significantly from bacterial cells. Therefore, optimizing these FPs to eukaryotic systems would require adaptations such as codon usage optimization for efficient expression, as well as potential adjustments to accommodate eukaryotic post-translational modifications and cellular environments.

## Methods

### General procedures

*E. coli* TB28 (MG1655 Δ*lacIZYA*) (Bernhardt and De Boer, 2003) was used all experiments except those involving ClpP fusions, which were performed in strain LY3581 (MG1655 Δ*clpP::FRT*).Strain LY3581 was constructed by P1 transduction of the *clpP* deletion from strain JW0427 (Baba et al., 2006). All experiments were performed by growing cells in M9 medium (referred to as M9GC): 7 g/L Na_2_HPO_4_.7H_2_O, 3 g/L KH_2_PO_4_, 0.5 g/L NaCl, 1 g/L NH_4_Cl, 1 mM MgSO_4_, 80 µM CaCl_2_ supplemented with 0.2% glucose, 0.4% casamino acids and 0.4 mg/L thiamine hydrochloride. Overnight cultures were inoculated in M9GC containing appropriate antibiotics (10 µg/mL chloramphenicol for pFN01 or 25 µg/mL kanamycin for pCLP) from colonies, followed by 1000x dilution in fresh M9GC medium without antibiotics. Cells were then grown to exponential phase, to a turbidity (OD_600_) of 0.2-0.3 at which experiments were performed. Fluorescence in bulk cultures was measured in 96 well plates.

All fluorescence measurements were performed on a Spark plate reader (Tecan) using a 5 nm bandpass monochromator. Cyan fluorescence was excited at 450 nm and collected at 480 nm, green fluorescence was excited at 480 nm and collected at 510 nm, yellow fluorescence was excited at 510 nm and collected at 540 nm, and red fluorescence was excited at 565 nm and collected at 595 nm.

### Molecular cloning

All plasmid constructs were obtained by Gibson assembly unless specified otherwise and are detailed in **Supplementary Table 1**. All primers used to construct these vectors are detailed in **Supplementary Table 2**. All enzymes were purchased from New England Biolabs. Polymerase chain reactions were performed using PrimeSTAR Max (Takara).Synthetic genes for mApple with modified N and C-termini and mGreenLantern were ordered from Integrated DNA Technologies.

pNF02 vectors used for unfused FP expression were constructed by amplifying a mini-F backbone from pNF02-mScarlet-I (Goormaghtigh et al., 2018) using primers bbpNF02 F & R.sfGFP was introduced in pNF02 using primers sfGFP02 F & R while mApple with new N and C-termini (mAppleNC) was directly inserted as a synthetic gene. Mutations to construct mChartreuse were introduced in pNF02-sfGFP using primers N39I F & 39 R, D129G F & I128S, N149K F & Y145F R, and V206K F & 206 R. mJuniper was generated from pNF02-mChartreuse using primers S72A F & Y66W, and H148D F & N146F R while mLemon was constructed using primers S72A F &S65GT63S F, V206K F & T203Y, and V224L F & 221 R. mLychee was obtained from pNF02-mAppleNC using primers V71A F & L85Q R,S131P F & R, A145P F & K139R R, and 219 F & I210V R.

pFN01 vectors, which encode all tested FPs fused to a reference FP and separated with a rigid linker, were constructed by amplifying a mTurquoise2-linker-mScarlet-I3 fragment from pDress-mTq2-link-mSc-I3 (Gadella et al., 2023) with primers IpFN01 F & R and a mini-F backbone from pNF02 using VpFN01 F & R. Red and yellow FPs were amplified using primer pairs 01miscF & R (mCherry, mApple,SYFP2), 01YPet F & R (mYPet), and 01sfGFP F & R (mLemon) on their respective templates (**Supplementary Table 1**) (Gadella et al., 2023;Shaner et al., 2008). To clone green and cyan FPs, mTurquoise2 was first replaced by mCherry on pFN01 by amplification of the backbone using VmTq2mCh F & R and by amplification of mCherry from pROD62 using ImTq2mCh F & R.Subsequently, FPs were cloned in this vector using primers 01sfGFP F & R (sfGFP, mChartreuse, mJuniper), 01mNGb F & R (mNeonGreen), mGL F & 01sfGFP R (mGreenLantern), and 01misc F & R (mTurquoise2) amplified from their respective templates (**Supplementary Table 1**) (Gadella et al., 2023; Rousseau et al., 2023). A pFN01-mCh-SCFP3A was generated by site-directed mutagenesis from pFN01-mCh-mTq2 using VSCFP3A F & R and ISCFP3A F & R.

The pCLP vector used to fuse FPs to ClpP was constructed by amplifying *clpP* and its native promoter using primers clpP F & R. This fragment was digested with NheI and BamHI and cloned in pUA66 (Zaslaver et al., 2006) digested with AvrII and BamHI. To construct ClpP-FP fusions, FPs were amplified from corresponding pFN01 vectors with primers insCLP F & R and cloned in a pCLP backbone amplified with pCLP F & R. Mutation S131P was introduced on pCLP-mCherry and pCLP-mApple using primers mFruitS131P F & R. pET151 vectors used for untagged FP production were constructed by amplification of a pET151 backbone using primers pET151 F & R, and of mChartreuse, mLemon, mJuniper or mLychee from pNF02 vectors using primers 151FP F & R.

### Maturation kinetics

TB28 cells carrying pFN01 vectors were grown to OD_600_ 0.2-0.3. A preheated 24-well plate was prepared with each well containing 500 µl M9GC medium supplemented with 200 µg/mL erythromycin, 20 µg/mL tetracycline hydrochloride & 20 µg/mL rifampicin. 500 µl of culture were mixed in each well, after which fluorescence acquisition was started with steps of 30 min for slow-maturing FPs (RFPs, mTurquoise2) or 5min for fast-maturing FPs with heating (37°C) and shaking every 5 min. A pseudo-first-order kinetic curve (F_(t)_=(F_max_-F_0_)*(1-e^-k*t^+F_0_) was fitted to each replicate using the least squares method, with F_max_ (fluorescence at plateau) & k (kinetic constant) as parameters, t (time) & F_(t)_ (fluorescence) as the variables, and F_0_ (fluorescence at time 0) being experimentally determined. Maturation half-time was calculated from these fitted kinetics as the time required to mature half of the remaining fluorescence signal (F_(t)_=(F_max_+F_0_)/2).

### Biophysical procedures

Untagged FPs were purified from BL21(DE3) cells carrying pET151 FP expression vectors. 50 ml cultures grown in LB (5 g/L yeast extract, 10 g/L tryptone, 5 g/L NaCl) at 30°C were brought to an OD_600_ of 0.6, after which FP production was induced overnight by the addition of 1 mM IPTG. Cells were subsequently pelleted and resuspended in 500 µl lysis buffer (50 mM Tris-Hcl pH 8.0, 300 mM NaCl, 5% glycerol and beaten with ∼300 µl 100 µm glass beads (5 min, 30 Hz) using a TissueLyser II (Qiagen). Lysates were separated from beads by piercing the bottom of the tube and by collecting the liquid phase in a bigger tube by centrifugation. After clearing by centrifugation, lysates are processed by three-phase partition (TPP) purification (Jain et al., 2004). A first TPP step was performed with 20% ammonium sulfate saturation & 1 volume t-butanol at room temperature. After phase separation by centrifugation, the aqueous phase of this TPP was subjected to a second step with 60% ammonium sulfate saturation & 2 volumes of t-butanol at room temperature, from which the interphase was isolated by centrifugation. This interphase was resuspended in 10 mM Tris-HCl pH 8.0 and ridden of insoluble impurities by centrifugation. A final desalting step was performed using Sephadex G-25 spin columns (Cytiva). Purified FPs were stored at 4°C in the dark and were left to mature at least 24 hours before experiments. Centrifugation of FP mother liquors was performed before all experiments to remove insoluble interfering impurities. All absorbance measurements were performed using FPs diluted in 10 mM Tris-HCl pH 8.0 to achieve a peak absorbance of 0.2, while all samples for fluorescence measurements were diluted to a peak absorbance of 0.04.

Extinction coefficients were determined as described (Cranfill et al., 2016), by normalizing the absorbance of FP samples to that of FPs denatured with 1 M NaOH, under the assumption that the molar extinction coefficient of the denatured chromophore at 447 nm (mChartreuse, mLemon) or 457 nm (mLychee) is 44 mM^-1^cm^-1^.Since mJuniper could not be denatured by strong bases, its concentration was determined by BCA assay (Pierce BCA Protein Assay Kit, Thermo Scientific). Quantum yields (QY) were determined as described (Cranfill et al., 2016), using fluorescein in 0.1 M NaOH (QY=0.85) (Berlman, 1971) as a reference for mChartreuse, mLemon and mJuniper, while rhodamine B in absolute ethanol (QY=0.49) (Casey and Quitevis, 1988) was used as a reference for mLychee.

Buffer titration to determine pKas was performed by mixing FP samples with 1 volume of citrate-phosphate-borate buffer at a given pH as described (Gadella et al., 2023). These buffers were prepared from a solution of 100 mM citric acid, 100 mM boric acid & 100 mM monosodium phosphate titrated with NaOH. Buffers were left to stabilize 24h at room temperature, yielding pH values of 3.01, 3.92, 4.92, 5.93, 7.07, 8.03, 8.90 & 9.94. Each replicate of fluorescence measurements was fitted to a Hill function (F_(pH)_=F_max_/(1+10^n*(pKa-pH^) using the least squares method, with F_(pH)_ (fluorescence at a given pH) and pH as variables, and F_max_ (fluorescence at plateau), n (Hill coefficient) and pKa as parameters.

### Microscopy procedures

Bacteria grown as described in general procedures were sealed on M9GC pads containing 1% agarose using GeneFrames (ThermoFisher). Cells were imaged using an Eclipse Ti2 microscope (Nikon) equipped with a Plan Apo λ 100x/1.45 objective, a motorized stage, Z-drift correction (Perfect Focus System, Nikon), heating (Okolab), a solid-state light source (Spectra X, Lumencor) and a sCMOS camera (Orca-Fusion BT, Hamamatsu). Cyan fluorescence was excited using a 438/24 excitation filter and collected using a 482/25 emission filter, green fluorescence was excited using a 474/27 excitation filter and a 515/30 emission filter, yellow fluorescence was excited using a 509/22 excitation filter and a 544/25 emission filter. Red fluorescent proteins were excited using a 578/21 excitation filter and a 641/75 emission filter for ClpP fusions or with a 553/24 excitation filter for photostability experiments. All fluorescent proteins were imaged using *ad-hoc* LEDs at 50% power intensity with 100 ms exposure times, except RFP photostability experiments which used 12% power of the 555 nm LED. Images were subsequently processed using MicrobeJ by automatic detection of cells and quantification of single-cell fluorescence parameters (*i*.*e*. mean for photostability experiments, skewness for ClpP fusions) (Ducret et al., 2016). Photobleaching experiments were performed by bleaching live cells with 100 ms steps using appropriate excitation settings as described above. Photobleaching half-times were determined as the time at which the mean fluorescence of the population dropped below 0.5.

### Resource availability

Expression plasmids for these new FPs were deposited on Addgene: pNF02-mChartreuse (219397), pNF02-mJuniper (219398), pNF02-mLemon (219399) & pNF02-mLychee (219400). Request for further information, data or materials should be addressed to the lead contact, C. Lesterlin.

## Acknowledgments

Plasmid pR6K-sfGFP was a kind gift from E. Gueguen. Plasmids pML31 & pROD62 were kind gifts from R. Reyes-Lamothe. Plasmids pDRESS-mTq2-link-mSc-I3 and pDX-mSc3-SYFP2 were kind gifts from T.W.J. Gadella obtained through Addgene (plasmids 189755 & 189764).Strain JW0427 was a kind gift from the National BioResource Project (NBRP, National Institute of Genetics, Japan). This work was supported by funding from the Foundation for Medical Research, Grant number FRM-EQU202103012587 to C.L. and A.C.) and from the French National Research Agency (Grant numbers ANR-22-CE12-0032 and ANR-23-CE12-0037 to C.L and N.F.).

## Author contributions

N.F. and C.L. conceptualized the paper. N.F. and A.C. performed experiments and analyzed the data.

N.F. wrote the paper and prepared the figures, with input from C.L. and A.C.. C.L. provided funding.

