## Supplementary Materials for "A palette of bright and photostable monomeric fluorescent proteins for bacterial time-lapse imaging"

| | 10 | 20 | 30 | 40 | |
|---|---|---|---|---|---|
| avGFP | M-SKGEELFT | GVVPILVELD | GDVNGHKFSV | SGEGEGDATY | GKLTLKFICT |
| mGreenlantern | MVSKGEELFT | GVVPILVELD | GDVNGHKFSV | RGEGEGDATN | GKLTLKFICT |
| sfGFP | M-SKGEELFT | GVVPILVELD | GDVNGHKFSV | RGEGEGDATN | GKLTLKFICT |
| mChatreuse | M-SKGEELFT | GVVPILVELD | GDVNGHKFSV | RGEGEGDAT**I** | GKLTLKFICT |
| mTurquoise2 | MVSKGEELFT | GVVPILVELD | GDVNGHKFSV | SGEGEGDATY | GKLTLKFICT |
| SCFP3A | MVSKGEELFT | GVVPILVELD | GDVNGHKFSV | SGEGEGDATY | GKLTLKFICT |
| mJuniper | M-SKGEELFT | GVVPILVELD | GDVNGHKFSV | RGEGEGDAT**I** | GKLTLKFICT |
| mYPet | M-SKGEELFT | GVVPILVELD | GDVNGHKFSV | SGEGEGDATY | GKLTLKLLCT |
| SYFP2 | MVSKGEELFT | GVVPILVELD | GDVNGHKFSV | SGEGEGDATY | GKLTLKLICT |
| mLemon | M-SKGEELFT | GVVPILVELD | GDVNGHKFSV | RGEGEGDAT**I** | GKLTLKFICT |

| | 50 | 60 | 70 | 80 | 90 |
|---|---|---|---|---|---|
| avGFP | TGKLPVPWPT | LVTTFSYGVQ | CFSRYPDHMK | QHDFFKSAMP | EGYVQERTIF |
| mGreenlantern | TGKLPVPWPT | LVTTLGYGVA | CFARYPDHMK | QHDFFKSAMP | EGYVQERTIS |
| sfGFP | TGKLPVPWPT | LVTTLTYGVQ | CFSRYPDHMK | QHDFFKSAMP | EGYVQERTIS |
| mChatreuse | TGKLPVPWPT | LVTTLTYGVQ | CFSRYPDHMK | QHDFFKSAMP | EGYVQERTIS |
| mTurquoise2 | TGKLPVPWPT | LVTTLSWGVQ | CFARYPDHMK | QHDFFKSAMP | EGYVQERTIF |
| SCFP3A | TGKLPVPWPT | LVTTLTWGVQ | CFARYPDHMK | QHDFFKSAMP | EGYVQERTIF |
| mJuniper | TGKLPVPWPT | LVTTLT**W**GVQ | CF**A**RYPDHMK | QHDFFKSAMP | EGYVQERTIS |
| mYPet | TGKLPVPWPT | LVTTLGYGVQ | CFARYPDHMK | QHDFFKSAMP | EGYVQERTIF |
| SYFP2 | TGKLPVPWPT | LVTTLGYGVQ | CFARYPDHMK | QHDFFKSAMP | EGYVQERTIF |
| mLemon | TGKLPVPWPT | LVT**S**L**G**YGVQ | CF**A**RYPDHMK | QHDFFKSAMP | EGYVQERTIS |

*** (above positions 65–67)

| | 100 | 110 | 120 | 130 | 140 |
|---|---|---|---|---|---|
| avGFP | FKDDGNYKTR | AEVKFEGDTL | VNRIELKGID | FKEDGNILGH | KLEYNYNSHN |
| mGreenlantern | FKDDGTYKTR | AEVKFEGDTL | VNRIVLKGID | FKEDGNILGH | KLEYNFNSHK |
| sfGFP | FKDDGTYKTR | AEVKFEGDTL | VNRIELKGID | FKEDGNILGH | KLEYNFNSHN |
| mChatreuse | FKDDGTYKTR | AEVKFEGDTL | VNRIELKG**SG** | FKEDGNILGH | KLEYN**Y**NSH**K** |
| mTurquoise2 | FKDDGNYKTR | AEVKFEGDTL | VNRIELKGID | FKEDGNILGH | KLEYNYFSDN |
| SCFP3A | FKDDGNYKTR | AEVKFEGDTL | VNRIELKGID | FKEDGNILGH | KLEYNYISDN |
| mJuniper | FKDDGTYKTR | AEVKFEGDTL | VNRIELKG**SG** | FKEDGNILGH | KLEYN**YF**S**DK** |
| mYPet | FKDDGNYKTR | AEVKFEGDTL | VNRIELKGID | FKEDGNILGH | KLEYNYNSHN |
| SYFP2 | FKDDGNYKTR | AEVKFEGDTL | VNRIELKGID | FKEDGNILGH | KLEYNYNSHN |
| mLemon | FKDDGTYKTR | AEVKFEGDTL | VNRIELKG**SG** | FKEDGNILGH | KLEYN**Y**NSH**K** |

| | 150 | 160 | 170 | 180 | 190 |
|---|---|---|---|---|---|
| avGFP | VYIMADKQKN | GIKVNFKIRH | NIEDGSVQLA | DHYQQNTPIG | DGPVLLPDNH |
| mGreenlantern | VYITADKQKN | GIKANFKTRH | NVEDGGVQLA | DHYQQNTPIG | DGPVLLPDNH |
| sfGFP | VYITADKQKN | GIKANFKIRH | NVEDGSVQLA | DHYQQNTPIG | DGPVLLPDNH |
| mChatreuse | VYITADKQKN | GIKANFKIRH | NVEDGSVQLA | DHYQQNTPIG | DGPVLLPDNH |
| mTurquoise2 | VYITADKQKN | GIKANFKIRH | NIEDGGVQLA | DHYQQNTPIG | DGPVLLPDNH |
| SCFP3A | VYITADKQKN | GIKANFKIRH | NIEDGGVQLA | DHYQQNTPIG | DGPVLLPDNH |
| mJuniper | VYITADKQKN | GIKANFKIRH | NVEDGSVQLA | DHYQQNTPIG | DGPVLLPDNH |
| mYPet | VYITADKQKN | GIKANFKIRH | NIEDGGVQLA | DHYQQNTPIG | DGPVLLPDNH |
| SYFP2 | VYITADKQKN | GIKANFKIRH | NIEDGGVQLA | DHYQQNTPIG | DGPVLLPDNH |
| mLemon | VYITADKQKN | GIKANFKIRH | NVEDGSVQLA | DHYQQNTPIG | DGPVLLPDNH |

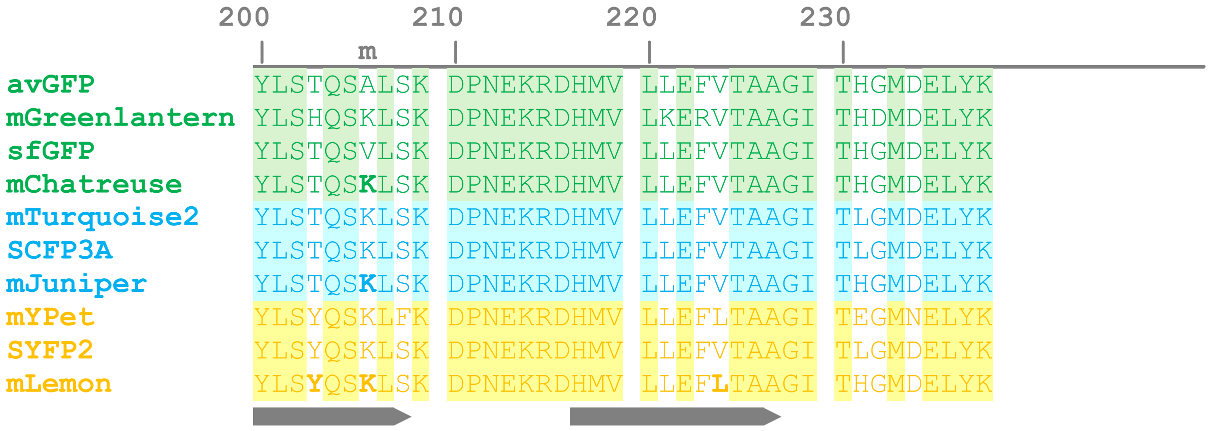


**Supplementary Figure 1: Alignments of avGFP derivatives used in this study.** Numbering is shown relative to avGFP. ɤ : alpha helixes ; ■► : beta sheets ; * : chromophore residues ; m : position of the monomerizing substitution (206K) (Zacharias et al., 2002). Substitutions introduced in the scope of this study are shown in bold. Note that our sfGFP variant does not contain the Q80R substitution as originally described and that our mYPet variant does not contain the valine insertion between positions 1 and 2. Also note that mNeongreen is not shown here due to low similarity with avGFP and derivatives.

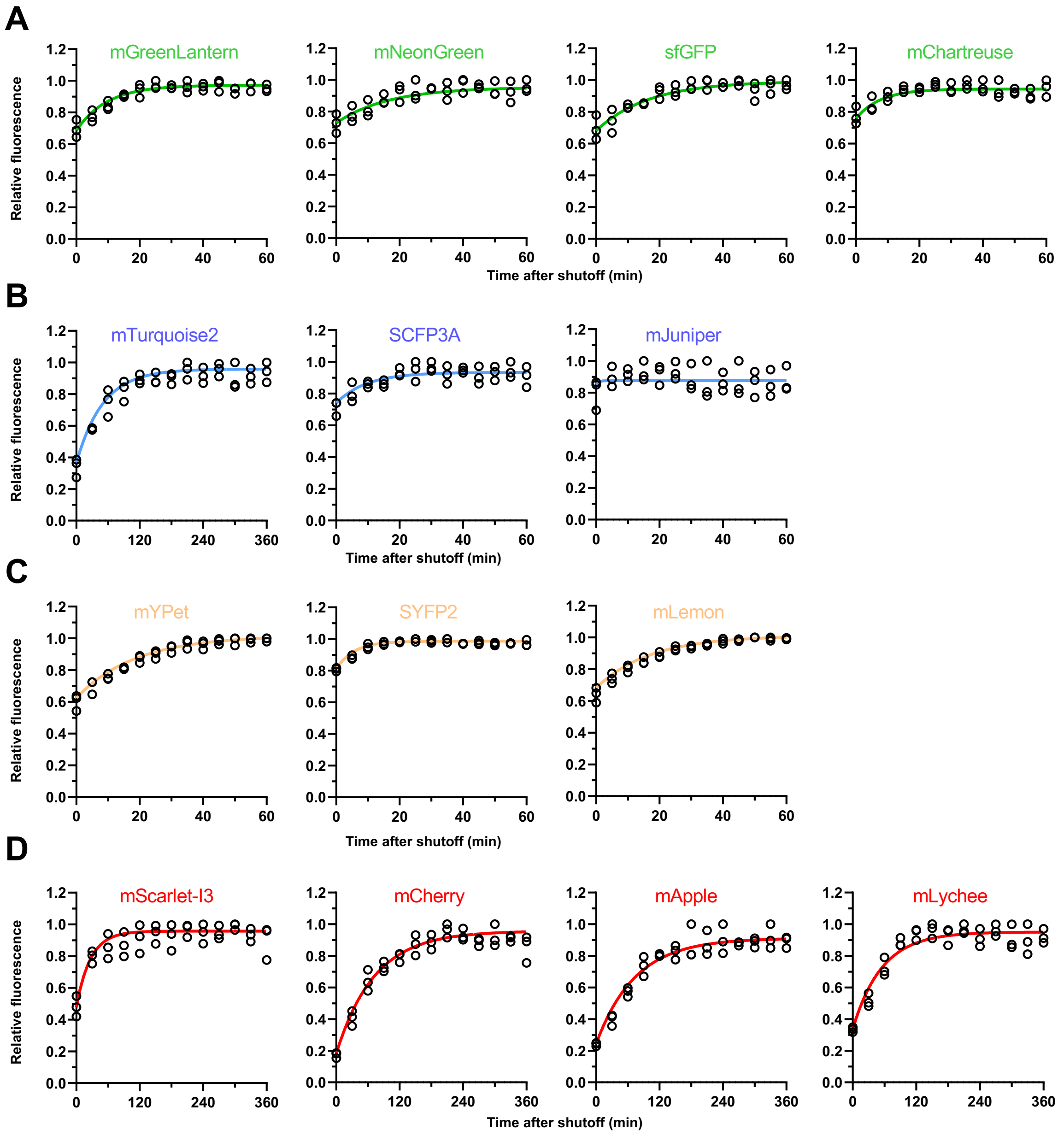


**Supplementary Figure 2: Maturation curves for green (A), cyan (B), yellow (C) and red (D) fluorescent proteins.** Exponentially-growing cultures of FP-producing bacteria were treated with 100 µg/mL erythromycin, 10 µg/mL tetracycline hydrochloride and 10 µg/mL rifampicin to arrest protein synthesis and enable dark fluorophores to fully mature. Fluorescence was read in 24-well plates every 5 min or 30 min under heating (37°C) and intermittent shaking. Data shows three independent replicates with a representative fit.

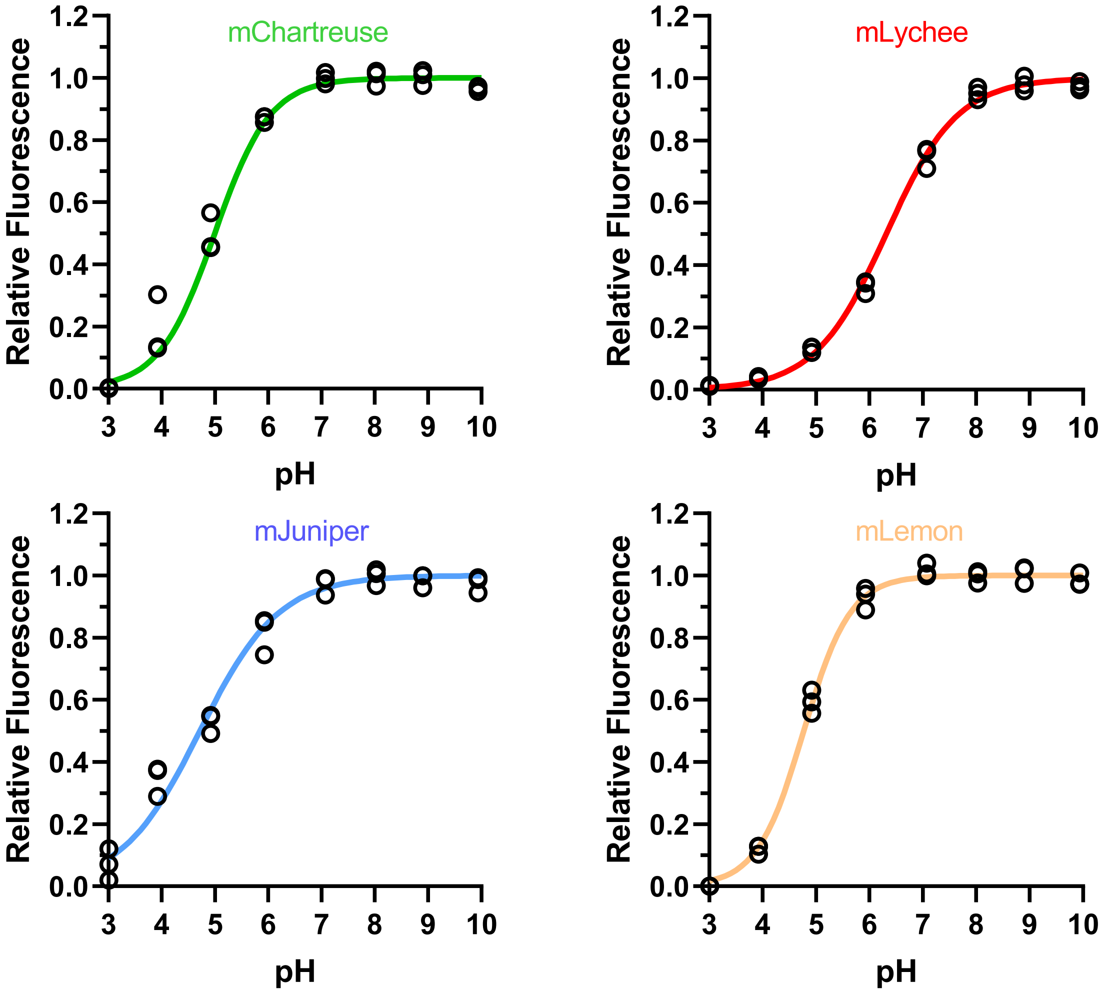


**Supplementary Figure 3: Effect of pH on fluorescence intensity.** Purified mChartreuse (A), mJuniper (B), mLemon (C) and mLychee (D) in 10 mM Tris-HCl pH 8.0 were diluted 2x in citrate-phosphate-borate buffers at indicated pH. Data shows the mean and standard deviation of three independent replicates with a fitted Hill function for each triplicate.

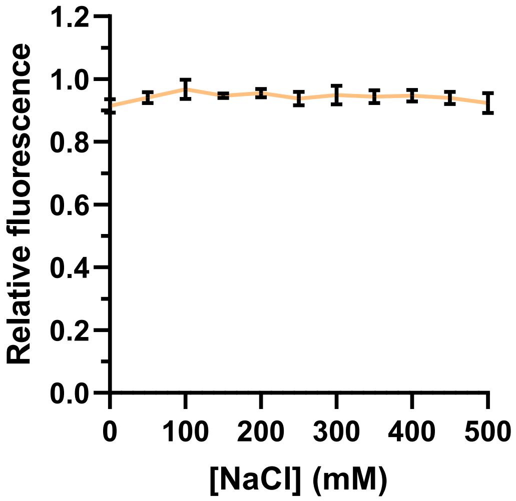


**Supplementary Figure 4 : Effect of sodium chloride on mLemon fluorescence intensity.** Purified mLemon in 10 mM Tris-HCl pH 8.0 was diluted in the same buffer containing increasing amounts of NaCl, with final concentrations shown on the x axis. Data shows three independent replicates with a representative fit.

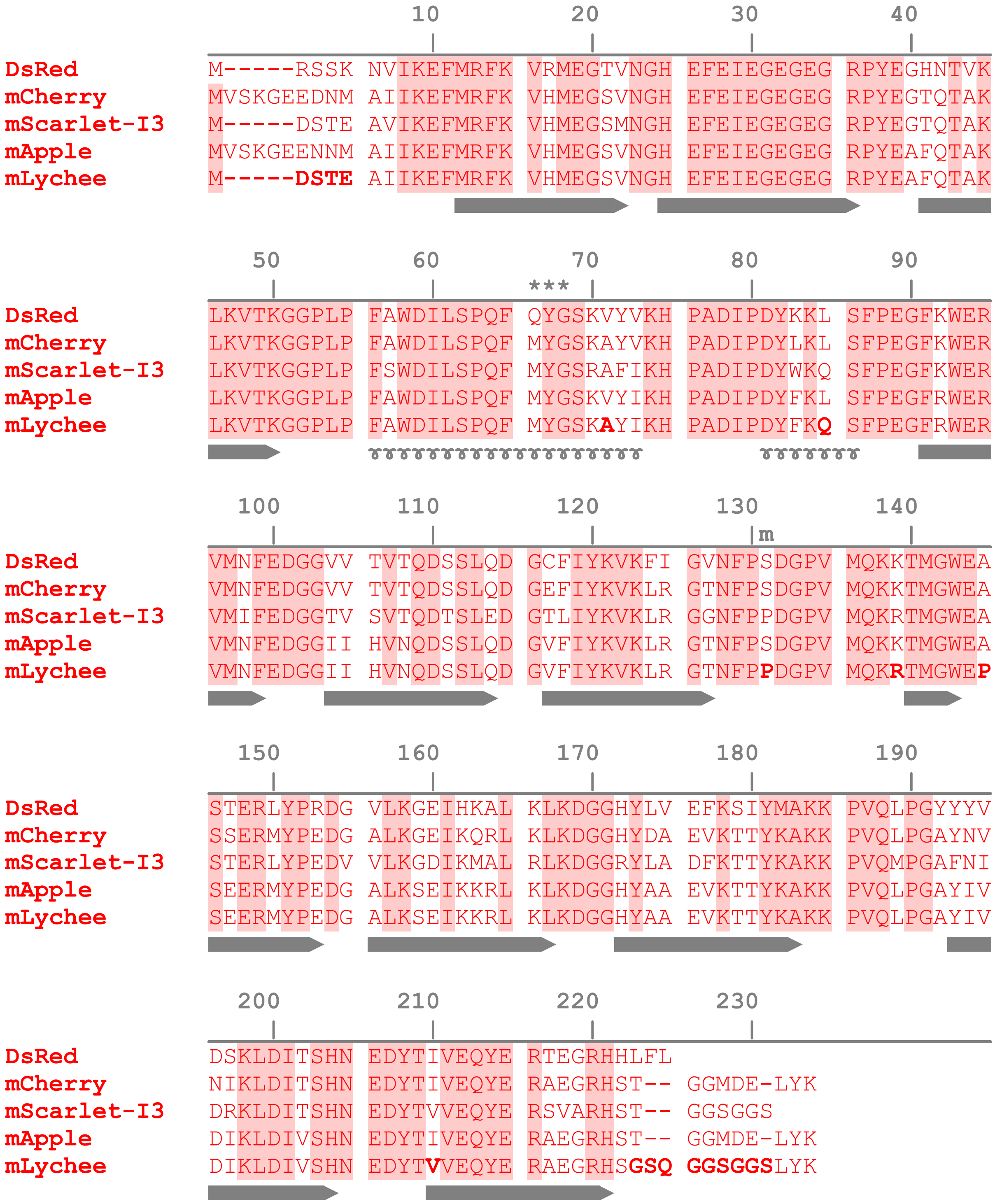


**Supplementary Figure 5 : Alignments of DsRed derivatives used in this study.** Numbering is shown relative to DsRed. ♉ : alpha helixes ; ➡ : beta sheets ; * : chromophore residues ; m : position of the monomerizing substitution (S131P) identified in this study. Substitutions introduced in the scope of this study are shown in bold.

**Supplementary Table 1 :** Plasmids & strains used in this study

| **Plasmid** | **Properties** | **Reference** |
|---|---|---|
| pNF02-mSc-I | Mini-F *cat proDp-mscarlet-I* | (Goormaghtigh et al., 2018) |
| pNF02-sfGFP | Mini-F *cat proDp-sfgfp* | This study |
| pNF02-mAppleNC | Mini-F *cat proDp-mapplenc* | This study |
| pNF02-mChartreuse | Mini-F *cat proDp-mchartreuse* | This study |
| pNF02-mJuniper | Mini-F *cat proDp-mjuniper* | This study |
| pNF02-mLemon | Mini-F *cat proDp-mlemon* | This study |
| pNF02-mLychee | Mini-F *cat proDp-mlychee* | This study |
| pDress-mTq2-link-mSc-I3 | Source for mTurquoise2 & mScarlet-I3 | (Gadella et al., 2023) |
| pR6K-sfGFP | Source for sfGFP | Lab collection, E. Gueguen |
| pNF02-mNG | Source for mNeongreen | (Rousseau et al., 2023) |
| pML31 | Source for mYPet | Lab collection, R. Reyes-Lamothe |
| pDx-mSc3-SYFP2 | Source for SYFP2 | (Gadella et al., 2023) |
| pROD62 | Source for mCherry | Lab collection, R. Reyes-Lamothe |
| mApple-pBAD | Source for mApple | (Shaner et al., 2008) |
| pFN01-mCh-sfGFP | Mini-F *cat proDp-mcherry-link-sfgfp* | This study |
| pFN01-mCh-mNG | Mini-F *cat proDp-mcherry-link-mneongreen* | This study |
| pFN01-mCh-mGL | Mini-F *cat proDp-mcherry-link-mgreenlantern* | This study |
| pFN01-mCh-mChartreuse | Mini-F *cat proDp-mcherry-link-mchartreuse* | This study |
| pFN01-mCh-mTq2 | Mini-F *cat proDp-mcherry-link-mturquoise2* | This study |
| pFN01-mCh-SCFP3A | Mini-F *cat proDp-mcherry-link-scfp3a* | This study |
| pFN01-mCh-mJuniper | Mini-F *cat proDp-mcherry-link-mjuniper* | This study |
| pFN01-mTq2-mYPet | Mini-F *cat proDp-mturquoise2-link-mypet* | This study |
| pFN01-mTq2-SYFP2 | Mini-F *cat proDp-mturquoise2-link-syfp2* | This study |
| pFN01-mTq2-mLemon | Mini-F *cat proDp-mturquoise2-link-mlemon* | This study |
| pFN01-mTq2-mCherry | Mini-F *cat proDp-mturquoise2-link-mcherry* | This study |
| pFN01-mTq2-mSc-I3 | Mini-F *cat proDp-mturquoise2-link-mscarlet-i3* | This study |
| pFN01-mTq2-mApple | Mini-F *cat proDp-mturquoise2-link-mapple* | This study |
| pFN01-mTq2-mLychee | Mini-F *cat proDp-mturquoise2-link-mlychee* | This study |
| pUA66 | $ori_{pSC101}$ *aphA gfpmut2* | (Zaslaver et al., 2006) |

| | | |
|---|---|---|
| pCLP | *ori*$_{pSC101}$ *aphA P*$_{clpXP}$*-clpP* | This study |
| pCLP-sfGFP | *ori*$_{pSC101}$ *aphA P*$_{clpXP}$*-clpP-sfgfp* | This study |
| pCLP-mNG | *ori*$_{pSC101}$ *aphA P*$_{clpXP}$*-clpP-mneongreen* | This study |
| pCLP-mGL | *ori*$_{pSC101}$ *aphA P*$_{clpXP}$*-clpP-mgreenlantern* | This study |
| pCLP-mChartreuse | *ori*$_{pSC101}$ *aphA P*$_{clpXP}$*-clpP-mchartreuse* | This study |
| pCLP-mTq2 | *ori*$_{pSC101}$ *aphA P*$_{clpXP}$*-clpP-mturquoise2* | This study |
| pCLP-SCFP3A | *ori*$_{pSC101}$ *aphA P*$_{clpXP}$*-clpP-scfp3a* | This study |
| pCLP-mJuniper | *ori*$_{pSC101}$ *aphA P*$_{clpXP}$*-clpP-mjuniper* | This study |
| pCLP-mYPet | *ori*$_{pSC101}$ *aphA P*$_{clpXP}$*-clpP-mypet* | This study |
| pCLP-SYFP2 | *ori*$_{pSC101}$ *aphA P*$_{clpXP}$*-clpP-syfp2* | This study |
| pCLP-mLemon | *ori*$_{pSC101}$ *aphA P*$_{clpXP}$*-clpP-mlemon* | This study |
| pCLP-mCherry | *ori*$_{pSC101}$ *aphA P*$_{clpXP}$*-clpP-mcherry* | This study |
| pCLP-mCherryS131P | *ori*$_{pSC101}$ *aphA P*$_{clpXP}$*-clpP-mcherry*$_{S131P}$ | This study |
| pCLP-mSc-I3 | *ori*$_{pSC101}$ *aphA P*$_{clpXP}$*-clpP-mscarlet-i3* | This study |
| pCLP-mApple | *ori*$_{pSC101}$ *aphA P*$_{clpXP}$*-clpP-mapple* | This study |
| pCLP-mAppleS131P | *ori*$_{pSC101}$ *aphA P*$_{clpXP}$*-clpP-mapple*$_{S131P}$ | This study |
| pCLP-mLychee | *ori*$_{pSC101}$ *aphA P*$_{clpXP}$*-clpP-mlychee* | This study |
| pCP20 | *ori*$_{pSC101(ts)}$ *bla cat cI857* P$_{L}$*-flp* | Lab collection |
| pET151 | *ori*$_{pBR322}$ *bla lacI* P$_{T7}$ | Life Technologies |
| pET151-mChartreuse | *ori*$_{pBR322}$ *bla lacI* P$_{T7}$*-mchartreuse* | This study |
| pET151-mJuniper | *ori*$_{pBR322}$ *bla lacI* P$_{T7}$*-mjuniper* | This study |
| pET151-mLemon | *ori*$_{pBR322}$ *bla lacI* P$_{T7}$*-mlemon* | This study |
| pET151-mLychee | *ori*$_{pBR322}$ *bla lacI* P$_{T7}$*-mlychee* | This study |
| **Strain** | **Genotype** | **Reference** |
| MG1655 | Wild-type *Escherichia coli* | Lab collection |
| TB28 | MG1655 Δ*lacIZYA::FRT* | (Bernhardt and De Boer, 2003) |
| JW0427 | BW25113 Δ*clpP::FRT-aphA-FRT* | (Baba et al., 2006) |
| LY3581 | MG1655 Δ*clpP::FRT* | This study |
| BL21(DE3) | Protein production *E. coli* strain | Lab collection |

**Supplementary Table 2 :** Primers used in this study

| Primer name | Primer sequence |
|---|---|
| bbpNF02 F | TAAGTGCACCTCTAGTATCACAC |
| bbpNF02 R | CATGCTAGCTTTCTCCTCTTTC |
| sfGFP02 F | GAAAGAGGAGAAAGCTAGCATGTCTAAAGGTGAAGAACTGTTC |
| sfGFP02 R | GTGATACTAGAGGTGCACTTATTTGTAGAGCTCATCCATGCCG |
| IpFN01 F | GGAGAAAGAAAAATGAAAACAGTGAGCAAGGGCGAGGAGC |
| IpFN01 R | CTAGAGGTGCACTTACTTGTACAAGGAGCC |
| VpFN01 F | CAAGTAAGTGCACCTCTAGTATCACAC |
| VpFN01 R | CACTGTTTTCATTTTTCTTTCTCCTCTTTCTCTAGTAAAAG |
| 01misc F | CCGGTCGCCACCATGGTGAGCAAGGGCGAGGAG |
| 01misc R | GTGTGATACTAGAGGTGCACTTACTTGTACAGCTCGTCCATGCC |
| 01YPet F | CCGGTCGCCACCATGTCTAAAGGTGAAGAATTATTCAC |
| 01YPet R | GTGTGATACTAGAGGTGCACTTATTTGTACAATTCATTCATACCCTC |
| 01sfGFP F | CCGGTCGCCACCATGTCTAAAGGTGAAGAACTG |
| 01sfGFP R | GTGTGATACTAGAGGTGCAC |
| VmTq2mCh F | GGCATGGACGAGCTCTACAAG |
| VmTq2mCh R | GCTCCTCGCCCTTGCTCAC |
| ImTq2mCh F | GTGAGCAAGGGCGAGGAGC |
| ImTq2mCh R | ACTTGTAGAGCTCGTCCATGCC |
| 01mNGb F | CCGGTCGCCACCATGGTTTCTAAAGGTGAAGAAGACAATATGGC |
| 01mNGb R | GTGTGATACTAGAGGTGCACTTATTTGTACAGTTCATCCATGCCC |
| mGL F | GCTACCGGTCGCCACCATGGTTTCTAAAGGTGAAG |
| VSCFP3A F | CAACTACATTAGCGACAACGTC |
| VSCFP3A R | CGCCCCAGGTCAGGGTGGTCACG |
| ISCFP3A F | GACCACCCTGACCTGGGGCGTGC |
| ISCFP3A R | CGTTGTCGCTAATGTAGTTGTACTCC |
| clpP F | CCCCGCTAGCGCTAAATTCGCACAAAGGC |
| clpP R | CCCGGATCCGCCACCGCCTGAATTACGATGGGTCAGAATCG |
| insCLP F | GTAATTCAGGCGGTGGCGGGCTACCGGTCGCCACCATG |
| insCLP R | CACGAGGCCCTTTCGTCTTGTGATACTAGAGGTGCACTTA |
| pCLP F | AAGACGAAAGGGCCTCGTG |
| pCLP R | CCGCCACCGCCTGAATTACG |
| mFruitS131P F | CCGACGGCCCCGTAATGCAG |
| mFruitS131P R | GGGGGAAGTTGGTGCCGCGC |
| pET151 F | CACCGCTGAGCAATAACTAGC |
| pET151 R | CTTTCTTAAAGTTAAACAAAATTATTTCTAGAGGGG |
| 151FP F | GAAATAATTTTGTTTAACTTTAAGAAAGAGGAGAAAGCTAGCATG |
| 151FP R | GTTATTGCTCAGCGGTGTGATACTAGAGGTGCAC |
